# From collocations to call-ocations: using linguistic methods to quantify animal call combinations

**DOI:** 10.1101/2021.06.16.448679

**Authors:** Alexandra B. Bosshard, Maël Leroux, Nicholas A. Lester, Balthasar Bickel, Sabine Stoll, Simon W. Townsend

## Abstract

Emerging data in a range of non-human animal species have highlighted a latent ability to combine certain pre-existing calls together into larger structures. Currently, however, there exists no objective quantification of call combinations. This is problematic because animal calls can co-occur with one another simply through chance alone. One common approach applied in language sciences to identify recurrent word combinations is collocation analysis. Through comparing the co-occurrence of two words with how each word combines with other words within a corpus, collocation analysis can highlight above chance, two-word combinations. Here, we demonstrate how this approach can also be applied to non-human animal communication systems by implementing it on a pseudo dataset. We argue collocation analysis represents a promising tool for identifying non-random, communicatively relevant call combinations in animals.

## Introduction

Over the last 20 years there has been growing interest into the combinatorial abilities of animals, namely the propensity to sequence context-specific calls (i.e. meaning-bearing units, see (Suzuki & Zuberbühler, 2019) into larger potentially meaningful structures (Arnold & Zuberbühler, 2006; Collier et al., 2020; Engesser, Ridley, & Townsend, 2016; Ouattara, Lemasson, & Zuberbühler, 2009; Suzuki, Wheatcroft, & Griesser, 2016). Combinatoriality is one mechanism that can increase the expressive potential of a finite vocal repertoire. It therefore provides important comparative insights into the complexity of animal vocal systems and the selective pressures such systems have been exposed to (Collier et al., 2020). These data also hold great promise in furthering our understanding of the similarities between animal communication and human language given that, for many years, it was assumed that the systematic concatenation of meaning-bearing units (i.e., syntax) was a phenomenon unique to language (Hurford, 2012). Emerging examples of syntactic-like structure in non-human primates and non-primate animals suggests this particular assumption was indeed premature (Arnold & Zuberbühler, 2006; Berthet et al., 2019; Coye, Ouattara, Zuberbühler, & Lemasson, 2015; Coye, Zuberbühler, & Lemasson, 2016) and such data even have the potential to further our understanding of the evolutionary progression of our own communication system (Leroux & Townsend, 2020; Townsend, Engesser, Stoll, Zuberbühler, & Bickel, 2018).

In light of the communicative and evolutionary insights that research on combinatoriality can provide, it is surprising that, to date, no objective means of quantifying call combinations has been proposed. This is problematic as animal calls may occur in rapid succession through chance alone, representing mere read-outs of contextual shifts. A method to capture greater-than-chance co-occurrence of calls is therefore central to reliably detecting and identifying non-random (i.e., potentially communicatively relevant) animal call combinations.

Similar methodological issues have been encountered in research on language learning and use (Bartsch, 2004; Evert, 2008; Gablasova, Brezina, & McEnery, 2017; Gries, 2013). One approach frequently implemented to identify combinations of words (mostly bigrams i.e., two-word/two-call structures) in large written and spoken corpora is collocation analysis (for a review, see Gries, 2013). Collocation analyses can take several forms, but the core commonality is that it contrasts the frequency with which specific words combine to measure the relative exclusivity of their relationship within a corpus (Church, Gale, Hanks, & Hindle, 1991; Gries & Stefanowitsch, 2004; Kennedy, 1991; Nesselhauf, 2005). In other words, such analyses reveal whether particular word/call combinations are more common than would be expected given an assumed random baseline (e.g., the uniform distribution, in which each combination is equally likely). For example, in English “*drink*” collocates with “*coffee*” and “*going*” collocates with “*to*” (to form the future tense or describe a motion event). Thus, collocation analyses can be understood as statistical measures of the influence that a lexical item has on its neighbours. It has since become a crucial tool in Corpus Linguistics for analysing lexical items and grammatical features of natural language (e.g., Bartsch, 2004; Lehecka, 2015; Stefanowitsch & Gries, 2003; Xiao & Mcenery, 2006).

In this paper we propose that by considering animal vocal data in a similar way as to how language data are treated (i.e., as a corpus) affords the unique opportunity to apply a variety of analytical tools habitually implemented in language sciences to study similar questions. Specifically, we demonstrate the application of collocation analyses to non-human animal datasets as a way to empirically identify combinations of two calls, henceforth termed *bigrams,* and the relative merits of doing so. We focus on two specific forms of collocation analysis commonly implemented in language sciences: Multiple Distinctive and Mutual Information Collocation Analyses (Gries, 2014).

### Collocation analyses - Multiple Distinctive and Mutual Information approaches

Multiple Distinctive Collocation Analysis (MDCA) is primarily used when investigating and testing for the degree of attraction between meaning-bearing units that share semantic similarities in grammatical constructions. Specifically, MDCA statistically contrasts all possible bigram combinations to estimate whether a given bigram occurs at frequencies higher or lower than what would be expected by chance (Gries & Stefanowitsch, 2004; Hilpert, 2006). Furthermore, the output of MDCA also provides a superficial estimate of bigram ordering, namely whether the combination is sensitive to the position of the calls comprising it (e.g., is A-B as frequent as B-A). Importantly, MDCA is not constrained by the usual sampling assumptions, making them suitable for skewed, non-random, and small corpora of the kind we tend to have in animal communication (Gries, 2014; Gries & Stefanowitsch, 2004; Hilpert, 2006).

One recurrent issue for the analysis of linguistic corpora is the fact that any corpus represents an incomplete – or undersampled – representation of the target linguistic system – i.e., some two-word combinations can be under-represented or even absent from a corpus when their “true” probability of occurrence is higher (note that this also affects all other probabilities in the sample, which will be artificially inflated). Such undersampling leads to misleading estimates of the significance of certain bigrams in the corpus (Gries & Stefanowitsch, 2004; Hilpert, 2006). Mutual Information Collocation Analyses (MICA) however actually overestimates to low-frequency values (Church & Hanks, 1990; Evert, 2005), which can be an advantage in animal communication corpora, as low-frequency pairings, a common feature of non-human vocal data sets, are not overlooked but flagged as potential combinatorial candidates. MICA calculates the variability of co-occurring items through computing information values via observed frequency divided by expected frequency. MICA can therefore – to some degree – better account for low-frequency bigrams, a common occurrence in small corpora and particularly in non-human vocal datasets.

In the remainder of the paper, we apply both forms of collocation analyses using an existing R script provided by Stefan Gries (2014, see supplementary) to a pseudo data set we built for the “Yeti” - a mythical ape-like creature.

#### Call combinations in Yetis: an example

We created an artificial set of call combinations produced by the Yeti (see Table 1). The data set was generated with a range of distributions in mind: specifically call combinations with i) high-frequencies, ii) low frequencies, and call types that appear in combination with a) only one other call type or b) with many different call types. The repertoire and sample size were simulated consistent with the known communicative repertoires of other primates (Leroux et al., in press). Yetis have a call repertoire comprising 10 calls, ranging from tonal whistles and twitters to noisy barks and coughs. Initial assessment of the data suggested that some of the calls co-occur with others from the repertoire and some co-occurrences are more frequent than others. We therefore expected the collocation analyses to reveal at least some significant associations between calls. To test this prediction, we applied both the MDCA and MICA to our data set.

**Table 1:**
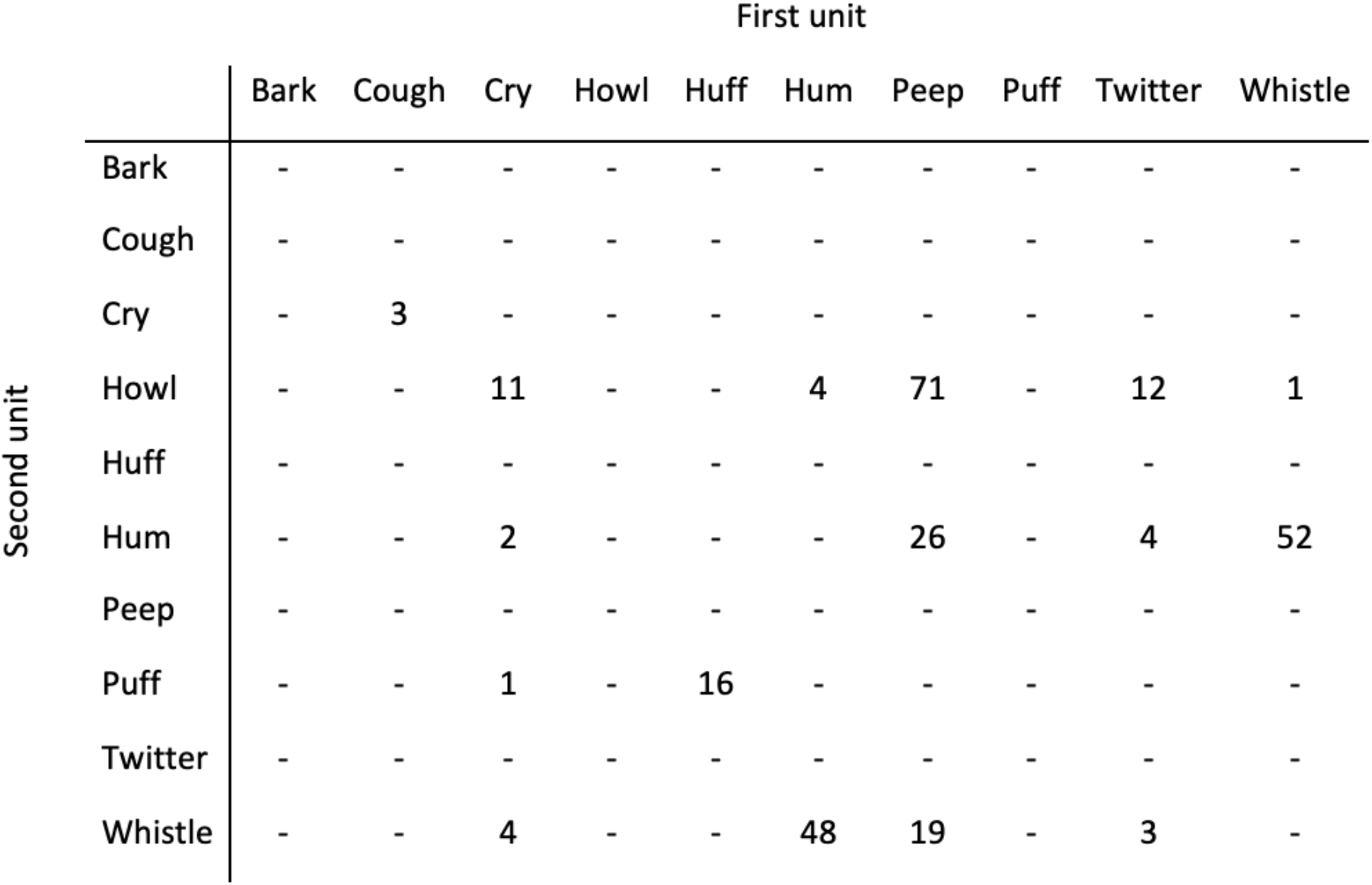
Distribution of bigrams occurring in the Yeti vocal repertoire. Columns and rows show the first and second unit within a call combination respectively.

#### Multiple Distinctive Collocation Analysis

Sixteen different potential bigrams were identified in the Yeti repertoire (see Table 1). In a first step we applied a Multiple Distinctive Collocation Analysis where call dependencies within these bigrams were calculated using an exact binomial test on each possible bigram combination (Gries, 2014; Gries & Stefanowitsch, 2004; Hilpert, 2006). Specifically, by applying a logarithmic transformation to p-values (used here explicitly since they reflect the relative association of calls but whilst simultaneously accounting for sample size), it is then possible to estimate whether a given bigram occurs at frequencies higher (positive “pbins”, Table 2) or lower (negative “pbins”, Table 2) than what would be expected by chance (i.e. the absolute value of pbin >3: P<0.001, *>2: P<0.01, *>1.3: P<0.05). Since the aim here is to identify potential candidates for meaningful call combinations, we will focus only on positive values that highlight an attraction between two call types. For positive pbin values, the higher the value for two calls, the greater their collocational strength.

**Table 2:** Multiple Distinctive Collocation analysis for the bigrams in the Yeti’s vocal data set. Columns and rows show the first and second unit within a call combination respectively. Values are pbins and can be translated to p-values (abs(pbin) *>3: P<0.001, *>2: P<0.01, *>1.3: P<0.05). Significant results are coloured in green.

|  |  | First unit |  |  |  |  |  |  |  |  |  |
| --- | --- | --- | --- | --- | --- | --- | --- | --- | --- | --- | --- |
|  |  | Bark | Cough | Cry | Howl | Huff | Hum | Peep | Puff | Twitter | Whistle |
| Second unit | Bark | - | - | - | - | - | - | - | - | - | - |
|  | Cough | - | - | - | - | - | - | - | - | - | - |
|  | Cry | - | 5.9 | -0.1 | - | -0.1 | -0.3 | -0.7 | - | -0.1 | -0.3 |
|  | Howl | - | -0.5 | 1.2 | - | -2.6 | -4.8 | 8.8 | - | 1.4 | -7.7 |
|  | Huff | - | - | - | - | - | - | - | - | - | - |
|  | Hum | - | -0.4 | -1.1 | - | -2.2 | -7.6 | -1.6 | - | -0.5 | 17.0 |
|  | Peep | - | - | - | - | - | - | - | - | - | - |
|  | Puff | - | -0.1 | -0.2 | - | 18.6 | -1.5 | -4.0 | - | -0.5 | -1.6 |
|  | Twitter | - | - | - | - | - | - | - | - | - | - |
|  | Whistle | - | -0.4 | -0.3 | - | -1.9 | 17.3 | -2.6 | - | -0.6 | -6.8 |

The Multiple Distinctive Collocation Analysis applied here suggests a significant relative attraction exists within six bigrams (see Table 2). The highest value was calculated for Huff-Puff, followed by Hum-Whistle, Whistle-Hum, Peep-Howl, Cough-Cry and lastly Twitter-Howl. Importantly, four of the six bigrams showed significant attraction between the two comprising calls in one specific order only (Huff-Puff, Peep-Howl, Cough-Cry & Twitter-Howl, Table 2). However, the combinations involving Hums and Whistles displayed significant attraction no matter the linearisation (Hum-Whistle and Whistle-Hum), suggesting that, either order did not matter for this particular combination, or that Whistle and Hum form two significant, differently ordered bigrams.

In light of the additional aforementioned advantages associated with an information-based approach, we complemented the MDCA with a MICA.

#### Mutual Information Collocation Analysis

Similarly to MDCA, MICA calculates the collocational strength of each specific call type with every other call type it collocates with. To do so, the *joint observed frequency* of a specific bigram is divided by its *joint expected frequency* and then logarithmically transformed.

Concretely, the number of times the calls actually appear in combination is divided by the number of times the calls would appear in combination if every call was randomly distributed throughout the dataset. Once more, the higher the collocation value, the stronger the collocational strength between two units (again, we focus on positive values that indicate an attraction only). As with MDCA, pbins represent the logarithmically transformed p-values (i.e., the absolute value of pbin >3: P<0.001, *>2: P<0.01, *>1.3: P<0.05).

Not accounting for any specific ordering of the structures in the received input, the Mutual Information Collocation Analysis demonstrated a significant relative attraction within two bigrams (Cough-Cry & Huff-Puff, see Table 3). MICA only highlighted bigrams with call types that appear exclusively in combination with their collocational partner and not many other call types, while not rendering significant values for combinations of call types that appear with more than one other call type in the corpus (see Table 1). It is also noteworthy that the MICA provides a strong value for a very low-frequency pairing, namely Cough-Cry, which can only be found three times in the recorded repertoire.

**Table 3:** Mutual Information Collocation analysis for the bigrams in the Yeti’s vocal data set. Columns and rows show the first and second unit within a call combination respectively. Values are pbins and can be translated to p-values (pbin *>3: P<0.001, *>2: P<0.01, *>1.3: P<0.05). Significant results are coloured in green.

|  | Bark | Cough | Cry | Howl | Huff | Hum | Peep | Puff | Twitter | Whistle |
| --- | --- | --- | --- | --- | --- | --- | --- | --- | --- | --- |
| Bark |  |  |  |  |  |  |  |  |  |  |
| Cough | - |  |  |  |  |  |  |  |  |  |
| Cry | - | 3.7 |  |  |  |  |  |  |  |  |
| Howl | - | - | 0.6 |  |  |  |  |  |  |  |
| Huff | - | - | - | - |  |  |  |  |  |  |
| Hum | - | - | -2.4 | - | - |  |  |  |  |  |
| Peep | - | - | - | 0.8 | - | -1.1 |  |  |  |  |
| Puff | - | - | -0.4 | - | 4.0 | - | - |  |  |  |
| Twitter | - | - | - | 0.8 | - | -1.2 | - | - |  |  |
| Whistle | - | - | -1.3 | - | - | 0.7 | -1.5 | - | -1.5 |  |

## Discussion

Here, we show that, when conceptualising animal vocal data in the same way as a language corpus, methods habitually implemented in corpus linguistics can be transferred reliably to non-human communication systems to highlight promising call combinations. We argue that this approach therefore represents a novel application of a more objective method to quantify the combinatorial dynamics of animal communication systems. Specifically, collocation analyses help disentangle “true”, or non-random, call combinations from happenstance juxtapositions of single calls. This is critical when investigating potentially meaningful structuring within animal vocal communication systems.

It is important to note that other systematic approaches to capture the sequential dynamics of animal vocal sequences (e.g., song), have been applied, including different Markovian and non-Markovian chain modelling (Kershenbaum et al., 2014; Sainburg, Theilman, Thielk, & Gentner, 2019; Suzuki, Buck, & Tyack, 2006). Nevertheless, we argue that there are strong advantages for using collocation analysis, since collocations are easily operationalisable across systems and provide a convenient and more descriptive (as opposed to modelling-based) account of the combinatorial dependencies between vocal units (Evert, 2005, 2008; Firth, 1957).

Whilst here we implement these analyses only with a pseudo data set, we have also applied MDCA and MICA in real-world settings, primarily when investigating the combinatorial properties of wild chimpanzee (Leroux et al., in press) and captive marmoset (Bosshard, 2020) vocal data sets. In these two primate examples, collocation analyses also reliably identified non-random combinations and, critically, these included ones we had already anecdotally highlighted as promising candidates in addition to others that were previously less clear to us during observational data collection.

Of particular relevance is the fact that the type of collocational analyses applied here were sensitive to bigrams even when they occurred very infrequently in the data set. This is because collocational analyses consider the exclusivity of the combinatorial relationship: if calls combine extremely rarely, they will still be detected as long as their relationship together is exclusive. Since considerable variation characterises the frequency of call combinations in animal communication (e.g., alarm call combinations are less frequent than social call combinations (Boesch & Crockford, 2005; Collier, Townsend, & Manser, 2017; Leroux, Chandia, Bosshard, Zuberbühler, & Townsend, in prep.), we can be confident that collocational analyses will identify all relevant combinations, both common and rare.

Another advantage of such collocation analyses is that they allow an estimate of the ordering of call combinations. Identifying variation in the temporal progression of calls is necessary to design experiments probing the role of order on meaning: data which are key to unpacking how similar animal call combinations and human language really are. Our results from the provided Yeti dataset indicate that some, but not all, of the identified combinations are characterised by ordering, again a finding that is replicated in our real-world data sets (Bosshard, 2020; Leroux et al., in press). This preliminary identification of call order therefore might serve as one possible additional filter when deciding which of the combinations detected from an animal data set to follow-up from an experimental/playback perspective.

In conclusion, we hope that the approach outlined here will be applied by other researchers in the field of animal communication as a way to disambiguate random from non-random combinatorial structures. Future work could also build on these initial approaches through applying other, as yet unexplored, sequence-based modelling methods currently used in language sciences to animal corpora (e.g., skip-gram modelling, see Guthrie, Allison, Liu, Guthrie, & Wilks, 2006). Ultimately, implementing the same objective, standardised methods such as that presented in this paper could allow researchers to make more meaningful comparisons both within and across systems, for example, at the individual, group, population or even species level.

## Acknowledgments

We thank members of the Comparative Communication and Cognition Group at the University of Zurich, specifically Piera Filippi, Mélissa Berthet, Joseph Mine, Anna J. Szmarowska and Manuel Rüdisühli for discussions. This work was supported by the Swiss National Science Foundation (PP00P3_163850) to S.W.T. and the NCCR Evolving Language (Swiss National Science Foundation Agreement #51NF40_180888). We declare no conflict of interest.

